# Detection of Influenza virus and *Streptococcus pneumoniae* in air sampled from co-infected ferrets and analysis of their influence on pathogen stability

**DOI:** 10.1101/2023.02.24.529988

**Authors:** Andrea J. French, Nicole C. Rockey, Valerie Le Sage, Karina Mueller Brown, Meredith J. Shephard, Sheila Frizzell, Mike M. Myerburg, N. Luisa Hiller, Seema S. Lakdawala

## Abstract

Secondary infection with *Streptococcus pneumoniae* has contributed significantly to morbidity and mortality during multiple influenza virus pandemics and remains a common threat today. During a concurrent infection, both pathogens can influence the transmission of each other, but the mechanisms behind this are unclear. In this study, condensation air sampling and cyclone bioaerosol sampling were performed using ferrets first infected with the 2009 H1N1 pandemic influenza virus (H1N1pdm09) and secondarily infected with *S. pneumoniae* strain D39 (Spn). We detected viable pathogens and microbial nucleic acid in expelled aerosols from co-infected ferrets, suggesting that these microbes could be present in the same respiratory expulsions. To assess whether microbial communities impact pathogen stability within an expelled droplet, we performed experiments measuring viral and bacterial persistence in 1 μL droplets. We observed that H1N1pdm09 stability was unchanged in the presence of Spn. Further, Spn stability was moderately increased in the presence of H1N1pdm09, although the degree of stabilization differed between airways surface liquid collected from individual patient cultures. These findings are the first to collect both pathogens from the air and in doing so, they provide insight into the interplay between these pathogens and their hosts.

**Importance:** The impact of microbial communities on transmission fitness and environmental persistence is under-studied. Environmental stability of microbes is crucial to identifying transmission risks and mitigation strategies, such as removal of contaminated aerosols and decontamination of surfaces. Co-infection with *S. pneumoniae* is very common during influenza virus infection, but little work has been done to understand whether *S. pneumoniae* alters stability of influenza virus, or vice versa, in a relevant system. Here, we demonstrate that influenza virus and *S. pneumoniae* are expelled by co-infected hosts. Our stability assays did not reveal any impact of *S. pneumoniae* on influenza virus stability, and a trend towards increased stability of *S. pneumoniae* in the presence of influenza viruses. Future work characterizing environmental persistence of viruses and bacteria should include microbially-complex solutions to better mimic physiologically relevant conditions.

## Observation and Discussion

Environmental stability of respiratory pathogens expelled from an infected host is a key factor impacting transmission (1). Previous work has shown that several factors (eg. humidity, temperature, and solute concentration) influence microbial stability in droplets (2–5). Our understanding of how microbes within the same droplets affect persistence is insufficient, as studies often only examine one microbe at a time. The limited work investigating how bacteria alter viral stability have primarily focused on enteric pathogen stability in feces and found that binding of poliovirus to bacteria increased virus stability (6–9). However, these studies did not examine how viruses alter bacterial stability. So, it remains unclear whether multiple microbes exist within the same aerosols, and if so, whether they influence each other to impact environmental persistence.

Bacterial co-infection is a common occurrence for viral respiratory pathogens: bacterial co-infection rates during influenza virus infection in humans range from 4.2-32.7% and cause significant illness in critically ill patients (10–12). Studies of influenza virus and *S. pneumoniae* secondary infection in animals have shown that influenza virus facilitates transmission of *S. pneumoniae* (13–15), while *S. pneumoniae* may decrease viral transmission (14, 15). Other groups have found that *S. pneumoniae* can increase influenza transmission after antibiotic administration (16). A study on the interaction of nasopharyngeal bacteria on influenza virus observed that influenza virus binds *S. pneumoniae* (17), suggesting that these pathogens may travel in the same aerosols. These observations indicate a complex interplay between these pathogens that requires further investigation to understand how their interactions affect environmental persistence and transmission.

### Co-infected ferrets shed H1N1pdm09 and Spn into expelled aerosols

Co-infections can lead to high titers of virus and bacteria in infected hosts (14, 15, 18), suggesting that multiple microbes could be present within expelled respiratory droplets. To characterize environmental shedding of H1N1pdm09 and Spn, ferrets were first infected with H1N1pdm09 and then infected with Spn 2 days later. Nasal washes were collected, and air sampling was performed for 3 days after co-infection.

Nasal wash titers from co-infected ferrets showed that all three animals shed H1N1pdm09 on days 3 and 4 post-H1N1pdm09 infection, but only two animals shed virus on day 5, while all animals shed Spn throughout the time course (Figure 1A). We next assessed whether infectious microbes were released from co-infected ferrets by air sampling with a condensation sampler (Supplemental Figure 1). Aerosolized infectious H1N1pdm09 was detected from all ferrets on day 3, but from fewer animals on days 4 and 5 (Figure 1B). Despite measurable levels of Spn in nasal washes, only one animal had viable Spn collected from the condensation sampler (Figure 1B). This may be underrepresenting expelled bacteria in the air, as previous work has shown that not all viable bacteria form colonies after aerosolization (19). Cyclone bioaerosol sampling, used to collect microbial genomic material, detected H1N1pdm09 in air samples from all co-infected ferrets for both the >4 μm and 1-4 μm fractions on all days (Figure 1C-D). The small <1 μm fraction had measurable H1N1pdm09 from two of three co-infected ferrets on any day (Figure 1E). Spn was only detectable in the >4 μm fraction in two animals (Figure 1C), which is unsurprising given that *S. pneumoniae* ranges from 5-10x greater in diameter than influenza and is, therefore, less likely to be found in smaller aerosols. This result may also underrepresent the amount of aerosolized Spn, since sample processing was not optimized for encapsulated bacterial DNA. Our results are the first to detect infectious H1N1pdm09 and viable Spn in expelled aerosols from co-infected animals.

**Figure 1.**
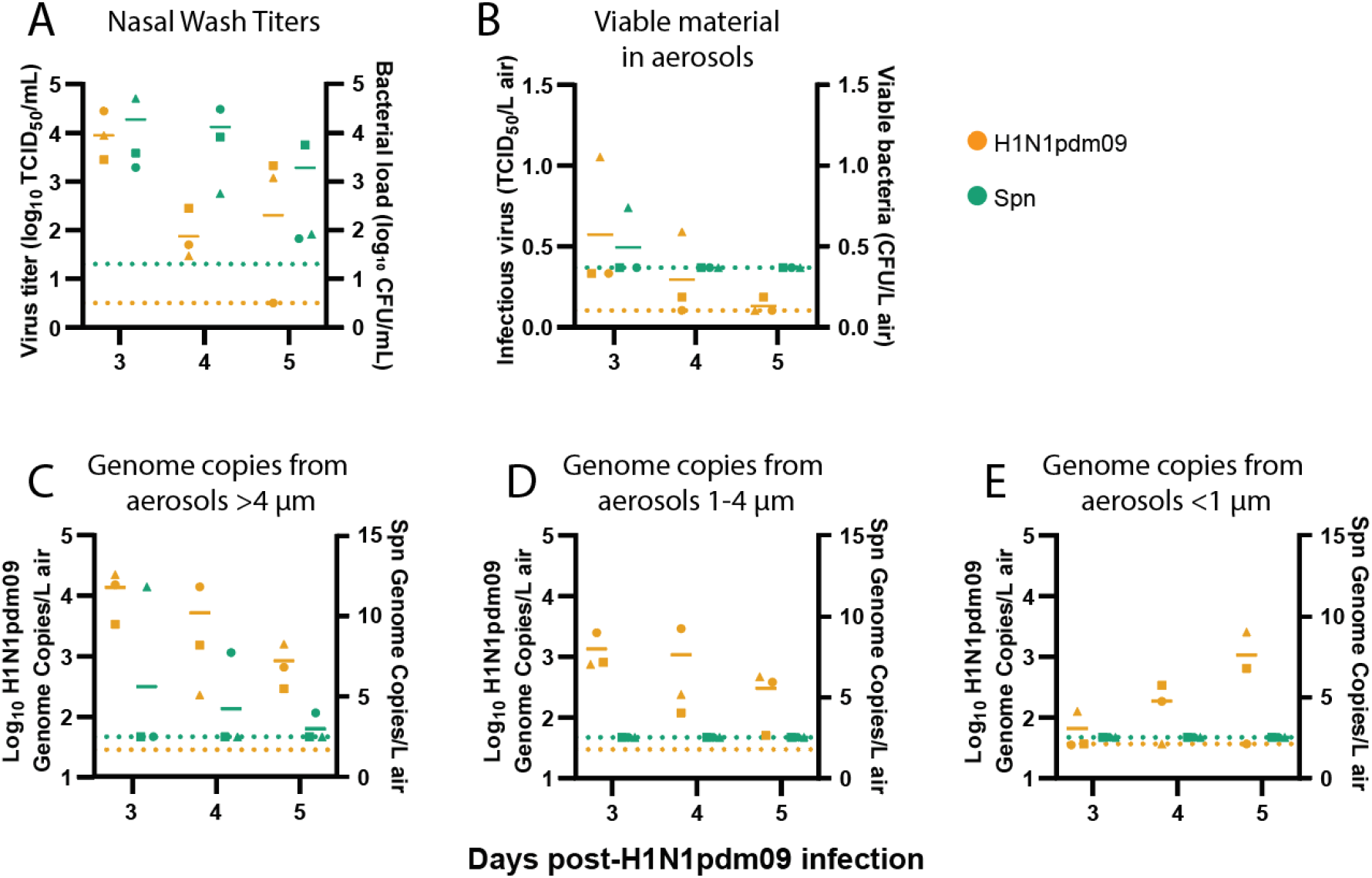
Co-infected ferrets shed H1N1pdm09 and Spn. Ferrets were infected with 10^6^ TCID_50_ of H1N1pdm09 and subsequently infected 2 days later with 10^7^ CFU *S. pneumoniae* D39. (**A**) Nasal wash loads of H1N1pdm09 and Spn are shown for the days following initial H1N1pdm09 infection. (**B**) Condensation sampling with a Liquid Spot Sampler was used to collect infectious virus and bacteria shed by co-infected animals. Viral and bacterial loads were measured by TCID_50_ and CFU assays, respectively. (**C-E**) Cyclone based air samplers were used to fractionate and collect microbial genomic material shed from co-infected ferrets in (**C**) >4 μm droplets, (**D**) 1-4 μm droplets, and (**E**) <1 μm droplets. Quantitative PCR was used to measure genome copies for each microbe. For all graphs, orange symbols represent H1N1pdm09 (N=3) and green symbols represent Spn (N=3), with each animal indicated by a unique shape and the mean indicated by short, solid lines. Dotted lines denote the limit of detection for H1N1pdm09 (orange) and Spn (green).

### Environmental stability of H1N1pdm09 is not impacted by the presence of Spn

Given the observation that H1N1pdm09 and Spn are shed from co-infected ferrets, we questioned whether these microbes might influence each other’s environmental stability in respiratory droplets. Spn has been shown to potentially alter influenza A transmission (14, 15), suggesting that Spn might decrease H1N1pdm09 environmental stability. H1N1pdm09, on the other hand, has been shown to increase transmission of Spn (13–15), which might indicate enhanced Spn stability with H1N1pdm09. To test this, we measured microbial persistence in droplets containing H1N1pdm09, Spn, or a 1:1 ratio of both pathogens in the presence of airway surface liquid (ASL) collected from four different human bronchial epithelial (HBE) cell donors (Figure 2). ASL is an important component of the respiratory tract and has been shown to increase stability of influenza viruses in the environment (3). After aging 1 μL droplets of each solution in a humidified chamber for 2 hours, infectious H1N1pdm09 or Spn was measured and compared to bulk solution controls (Figure 2A-B). Experiments were performed at 43% relative humidity (Figure 2E), as viruses and Gram-positive bacteria are more susceptible to decay at intermediate relative humidity (20). After 2 hours, there was no significant difference (p= 0.721) in H1N1pdm09 stability with or without Spn (average decay of 1.19 log_10_ TCID_50_/mL versus 1.34 log_10_ TCID_50_/mL, respectively) (Figure 2C). In contrast, there was a trend of increased stability for Spn in the presence of H1N1pdm09. Improved Spn stability was clearly observed in ASL from one patient culture (0284), as Spn alone decayed an average of 3.86 log_10_ CFU/mL and H1N1pdm09/Spn decayed an average of 2.81 log_10_ CFU/mL (p=0.096, Figure 2D, Supplemental Table 1). More modest stabilization for Spn was observed in 2 other cultures (0223 and 0259) and no difference was observed in ASL from one patient (0305) (Supplemental Table 1). Together, these results suggest that H1N1pdm09 infectivity is not impacted by Spn at the environmental conditions tested. There may be a modest impact of Spn stability in the presence of H1N1pdm09, although this may be more sensitive to variations in the ASL (or mucus composition) per individual. Further research should explore the impact of mucus and lung disease states on the relationship between influenza viruses and *S. pneumoniae*.

**Figure 2.**
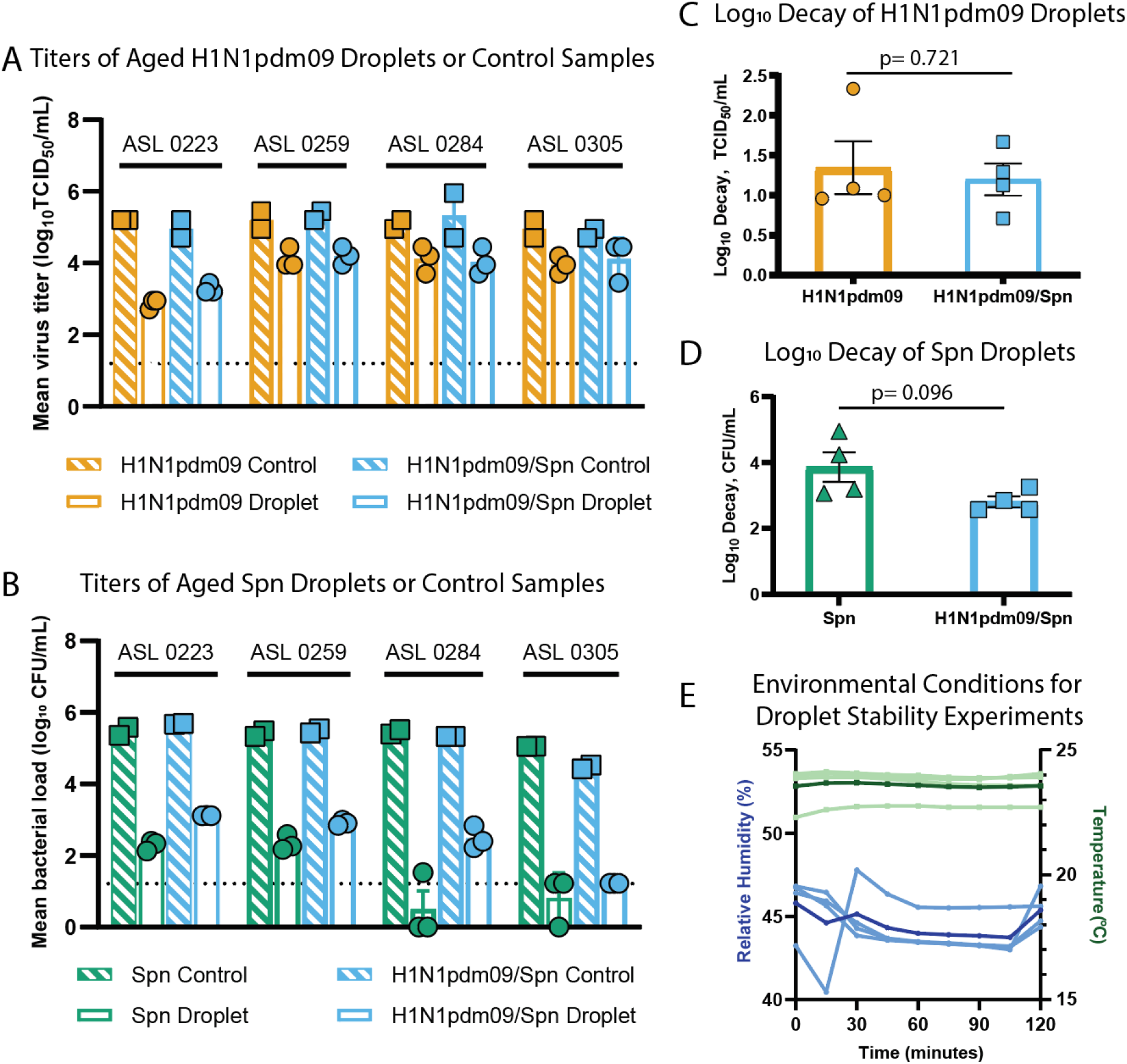
Stability of *S. pneumoniae* and influenza viruses in droplets. (**A-D**) Viral and bacterial loads of H1N1pdm09 and Spn were assessed after exposure of 10×1 μL droplets to 43% relative humidity (RH) at room temperature for 2 hours. Microbes were suspended in ASL from four different HBE cell donors as indicated in **A** and **B**. Control titers were determined using 10 μL of bulk solutions in closed tubes at room temperature. (**A**) The stability of H1N1pdm09 in droplets containing H1N1pdm09 or H1N1pdm09/Spn measured by TCID_50_ assay, and (**C**) log_10_ decay for each individual ASL culture were determined. (**B**) The stability of Spn in droplets containing Spn or H1N1pdm09/Spn measured by CFU assay, and (**D**) log_10_ decay for each individual ASL culture were determined. Differences were assessed using Welch’s unpaired t-test. (**E**) The RH and temperature were recorded every 15 minutes during stability experiments. Temperature (light green) and RH (light blue) for each ASL replicate are shown. The average temperature (dark green) and RH (dark blue) for all experiments are also included.

**Supplemental Table 1.** Average log decay for H1N1pdm09 or Spn in droplets

|  |  |  |  |  |  |
| --- | --- | --- | --- | --- | --- |
|  |  | Log<sub>10</sub> Decay H1N1pdm09 ± SEM |  | Log<sub>10</sub> Decay Spn ± SEM |  |
| ASL Donor | HBE Donor Condition | H1N1pdm09 | H1N1pdm09/Spn | Spn | H1N1pdm09/Spn |
| 0223 | COPD | 2.333 ± 0.068* | 1.667 ± 0.068* | 3.187 ± 0.067* | 2.565 ± 0.002* |
| 0259 | COPD | 1.083 ± 0.136 | 1.125 ± 0.118 | 3.071 ± 0.101* | 2.577 ± 0.033* |
| 0284 | IPF | 0.958 ± 0.180 | 1.292 ± 0.180 | 4.950 ± 0.415* | 2.848 ± 0.148* |
| 0305 | COPD | 1.00 ± 0.118 | 0.708 ± 0.272 | 4.237 ± 0.333 | 3.256 ± 0 |
| An asterisk indicates the p-value was less than 0.05 when comparing droplets of individual microbes to droplets with both microbes using a Welch's unpaired t-tests. |  |  |  |  |  |

Co-infection with pathogens can impact the transmissibility to subsequent hosts. Concurrent infections of influenza virus and *S. pneumoniae* result in increased morbidity and a greater risk of bacterial transmission (13, 14). The work here shows that co-infected animals expel both influenza virus and *S. pneumoniae* into air and can be collected using a condensation air-sampler or cyclone bioaerosol sampler. No impact was observed for influenza virus in the presence of *S. pneumoniae*, but a trend towards increased *S. pneumoniae* stability in the presence of influenza virus may help explain augmented *S. pneumoniae* transmission in addition to the increased bacterial shedding observed during co-infection (13, 14). Investigation of microbial stability using polymicrobial populations is not widely performed and could help elucidate the complexity of pathogen transmission seen in the human population. In addition, identifying host specific factors underlying microbial stability in the environment could increase our understanding of individual transmission risks and strategies mitigating the spread of pathogens in the population.

## Methods

Methods can be found in the supplemental materials.

## Acknowledgements

We thank members of the Lakdawala and Hiller lab for providing constructive feedback of this manuscript and Dr. Rachel Duron for editorial comments. This work was supported in part with Federal funds from NIAID, NIH, and DHHS (75N93021C00015, SSL). Additional funding was provided by Flu Lab (SSL and LCM) and NIAID (R01 AI158484). JF was supported by the University of Pittsburgh Training Program in Antimicrobial Resistance (T32AI138954). We would also like to acknowledge the Research Development Program from the Cystic Fibrosis Foundation to the University of Pittsburgh (MM and SF).

## Supplemental Materials and Methods

### Virus and Bacteria

A/California/07/2009 (H1N1pdm09) was grown in minimum essential media in Madin-Darby Canine Kidney (MDCK) cells at 37°C for 48 hours and collected by centrifuging supernatant to remove cell debris. Quantification of virus was performed using the 50% tissue culture infectious dose assay (TCID_50_) of 10-fold serial dilution on MDKC cells in 96-well plates and subsequent assessment for cytopathic effects 4 days after plating.

*S. pneumoniae* D39 (Spn) was grown in Columbia broth at 37°C. Quantification of bacterial burden was performed by plating 10-fold serial dilutions on blood agar plates and counting colony-forming units after incubation at 37°C overnight.

### Animals

Experiments involving ferrets were performed at the University of Pittsburgh under BSL2 safety conditions (IACUC protocol 19075697). Four to six-month male ferrets were confirmed to be seronegative for influenza infection prior to purchase. Animals were intranasally infected with 10^6^ TCID_50_ of H1N1pdm09 in 500 μL total volume and 10^7^ CFU of Spn in 500 μL. Ferrets were sedated using isoflurane prior to nasal wash collection, performed by collecting the flow-through of PBS passed through the nostrils.

### Air Sampling

Infectious virus and bacteria were collected using the Liquid Spot Sampler (Aerosol Devices Inc, Series 110), which uses condensation to collect aerosols into a collection vial. Air was collected from infected animals in a 7 liter chamber connected to the Spot sampler via anti-static tubing for 15 minutes each day at a rate of 1.4L/minute (Supplemental Figure 1). Sampling was performed on days 3, 4, and 5 post-H1N1pdm09 infection (days 1, 2, and 3 post-Spn infection) and prior to nasal wash collection. Condensed aerosols were collected in 700μL 0.5% BSA in PBS. Samples were immediately plated to quantify expelled bacteria and the remaining sample was used for virus titration as described above.

Aerosol sampling of H1N1pdm09/Spn-infected ferrets was performed using cyclone-based air samplers (BC251 developed by NIOSH) on days 3, 4, and 5 post-H1N1pdm09 infection to collect microbial genomic material. Samplers, calibrated to collect 3.5L of air per minute, were placed downwind of infected animals in cages with directional airflow and were run for 1 hour. Samplers fractionated aerosols into three sizes: aerosols >4μm, 1-4μm, and <1μm diameter. After aerosol collection, samplers were washed with isopropanol and allowed to air-dry to avoid contamination.

RNA was isolated using 500μL MagMAX Lysis/Binding Solution Concentrate in each collection tube with thorough vortexing. QIAamp viral RNA mini kit was used to isolate DNA and RNA from lysis solution. Viral and bacterial genome copies were quantified using RT-qPCR with primers against influenza M gene (Forward 5’-AGATGAGTCTTCTAACCGAGGTCG-3’; Reverse 5’-GCAAAGACACTTTCCAGTCTCTG-3’; Probe 5’- [FAM]TCAGGCCCCCTCAAAGCCGA[3BHQ1]-3’) or *S. pneumoniae* lytA gene (Forward 5’- ACGCAATCTAGCAGATGAAGCA-3’; Reverse 5’-TCGTGCGTTTTAATTCCAGCT-3’; Probe 5’-[HEX]GCCGAAAACGCTTGATACAGGGAG[BHQ1]-3’). *In vitro* transcribed RNA was used to make a standard curve for influenza virus, and *S. pneumoniae* genomic DNA was serially diluted to generate a standard curve for Spn. Limits of detection were determined by a Ct = 40 or a positive day 0 sample.

### Stability Experiments

Inside a biosafety cabinet, a saturated salt solution of K_2_CO_3_ was used to condition a glass chamber to 43% relative humidity, and a HOBO UX-100-011 logger was used to record temperature and humidity conditions during each ASL replicate (Figure 2E). Experimental solutions were generated using 10^7.15^ CFU/mL Spn, 10^7.15^ TCID_50_/mL H1N1pdm09, and a 1:5 dilution in PBS of airway surface liquid collected from human bronchial epithelial cells. Ten 1 μL droplets were incubated on polystyrene tissue culture plates in the conditioned chamber for two hours. Controls were 10 μL samples of each microbial solution in closed tubes that were incubated for 2 hours at ambient temperature during the chamber experiments. Log_10_ decay was calculated as previously described and represents the loss in virus or bacterial infectivity (21). Log_10_ decay was determined for each droplet replicate by subtracting the titer of the droplets from the average of the controls for the corresponding ASL. Experiments were performed using technical triplicates for droplets and technical duplicates for controls.

Human lung tissue collected using an approved protocol was used to differentiate human bronchial epithelial cells as previously described (22). Airway surface liquid was collected by washing differentiated cells with 150 μL PBS and collecting the wash (3). All HBE donors were diagnosed with chronic obstructive pulmonary disease (COPD), except for HBE 0284, which came from a patient diagnosed with idiopathic pulmonary fibrosis.

## Data Availability

The data that supports the findings shown here will be made openly available in FigShare at DOI: 10.6084/m9.figshare.22129055 upon publication. Some of the stability experiments were previously made available on BioRxiv at https://doi.org/10.1101/2020.11.10.376442

---

**Supplemental Figure 1.**
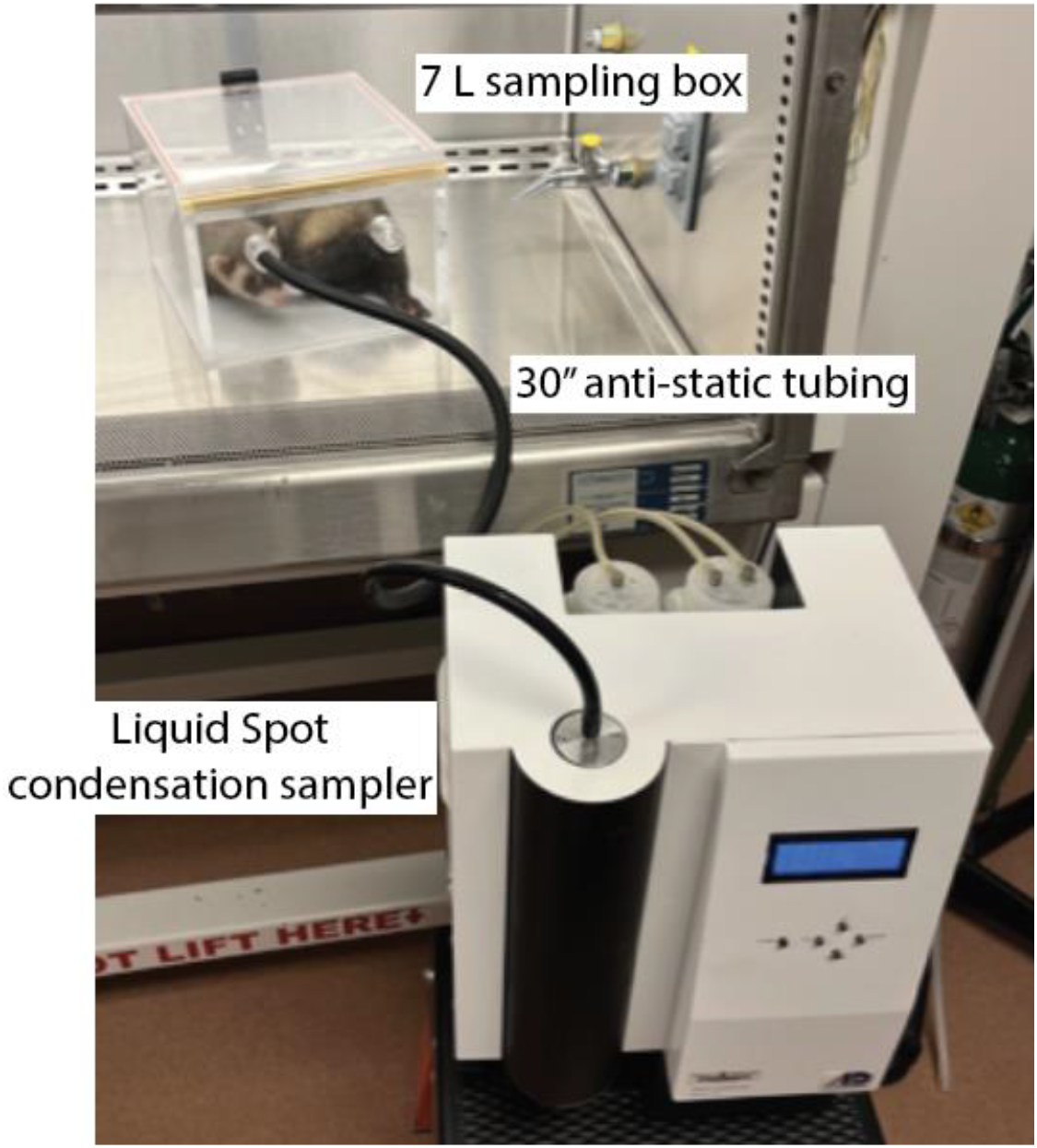
The Liquid Spot condensation sampler was used to collect infectious material from coinfected animals. Co-infected animals were placed in a sampling box for 15 minutes while sampling was performed. Anti-static tubing connected the sampling box to the inlet of the sampler.

